# ATOMIC: A graph attention neural network for ATOpic dermatitis prediction on human gut MICrobiome

**DOI:** 10.1101/2025.07.14.664646

**Authors:** Hyunsu Bong, Joonhong Min, Songhyeon Kim, Wootaek Lim, Dongyoung Lim, Hyunjeong Eom, Young Her, Minji Jeon

**Affiliations:** Department of Medicine, Korea University College of Medicine, Seoul, Korea; Department of Dermatology, Kangwon National University Hospital, Kangwon National University College of Medicine, Chuncheon, Korea; Woodang Network, Korea; Department of Dermatology, School of Medicine, Kangwon National University, Chuncheon, Korea; Department of Biomedical Informatics, Korea University College of Medicine, Seoul, Korea; Biomedical Research Center, Korea University Anam Hospital, Seoul, Korea

## Abstract

Atopic dermatitis (AD) is a chronic inflammatory skin disease driven by complex interactions among genetic, environmental, and microbial factors; however, its etiology remains unclear. Recent studies have reported the role of gut microbiota dysbiosis in AD pathogenesis, leading to increased interest in microbiome-targeted therapeutic strategies such as probiotics and fecal microbiota transplantation. Building on these findings, recent advances in computational modeling have introduced machine learning and deep learning-based approaches to capture the nonlinear relationships between gut microbiota and diseases. However, these models focus on diseases other than AD and often fail to capture complex microbial interactions or incorporate microbial genomic information, thereby offering limited interpretability. To address these limitations, we propose ATOMIC, an interpretable graph attention network-based model that incorporates microbial co-expression networks to predict AD. Microbial co-expression networks incorporate microbial genomic information as node features, thereby enhancing their ability to capture functionally relevant microbial patterns. To train and test our model, we collected and processed 99 gut microbiome samples from adult patients with AD and healthy controls at Kangwon National University Hospital (KNUH). As a result, ATOMIC achieved an AUROC of 0.810 and an AUPRC of 0.927 on the KNUH dataset. Furthermore, ATOMIC identified microbes potentially associated with AD prediction and proposed candidate microbial biomarkers that may inform future therapeutic strategies. To facilitate future research, we publicly released a gut microbial abundance dataset from KNUH. The source code and processed abundance data are available from ATOMIC GitHub repository at https://www.github.com/KU-MedAI/ATOMIC.

**Author summary:** Atopic dermatitis (AD) is a chronic inflammatory skin disease affecting approximately 120 million people worldwide, often leading to a reduced quality of life, including sleep disturbances and stress. Current treatment methods rarely achieve complete remission, and the precise cause of AD remains unclear. The association between imbalances in the gut microbiota and the pathogenesis of AD has highlighted the gut microbiota as a promising target for therapeutic strategies. Exploring the potential role of the gut microbiota in modulating host immune responses is likely to have a positive impact on future AD treatment. Consequently, the growing need to develop machine learning and deep learning-based predictive models using gut microbiome data. However, existing models often suffer from limited interpretability owing to their black-box nature and frequently overlook microbial interactions and genomic contexts. To address these limitations, we propose a novel model called ATOMIC, which integrates microbial co-expression networks and genomic information to predict AD. Our model achieved superior performance in terms of AUROC, AUPRC, and F1 scores compared with existing models and provided interpretable insights through an attention mechanism, thereby offering promising potential for identifying relevant biomarkers. Together, these results highlight ATOMIC’s potential for the accurate prediction of AD and for identifying biologically meaningful microbial signatures.

## Introduction

Atopic dermatitis (AD) is a chronic inflammatory skin disease characterized by intense itching and eczema [1]. Affecting over 120 million individuals globally, its prevalence continues to rise [2]. Given its chronic and relapsing nature, AD significantly impacts quality of life and is often accompanied by psychological symptoms such as stress and sleep disturbances [3, 4]. Multiple factors, including immune-related genetic and environmental factors, skin barrier dysfunction, and microbiota imbalance are known to contribute to its pathogenesis [5–8]. However, the precise etiology of AD remains unclear [9]. Current treatments, such as dupilumab and topical corticosteroids, can alleviate symptoms but rarely achieve complete remission [10], highlighting the need for a deeper understanding of the disease mechanism.

Among these factors, increasing attention has been directed toward the role of gut microbiota in the pathogenesis of AD [11]. The gut microbiota, comprising approximately 10–100 trillion microorganisms, plays a critical role in maturation of the immune system [12]. Through colonization resistance, a competitive process among microbes for nutrients and space, the microbiota maintains homeostasis and suppress pathogenic organisms [13]. Notably, a reduction in the risk of developing AD has been observed in infants with high levels of short-chain fatty acids (SCFA), such as butyrate and propionate [14]. Given that SCFAs support epithelial barrier integrity and modulate cytokine production and immune responses [15], the depletion of SCFA-producing microbes may weaken colonization resistance and contribute to the development of AD.

With growing evidence linking the gut microbiota to AD, interventions such as probiotics and fecal microbiota transplantation (FMT) are being actively explored as novel therapeutic strategies [16]. Probiotics, defined as live microorganisms that provide health benefits upon ingestion [17], have been shown to alleviate AD symptoms by restoring gut microbiota balance [18]. FMT involves the transfer of fecal microbiota from healthy donors to patients [19] and has the potential to suppress AD-related allergic responses and improve immune regulation [16, 20]. However, the effectiveness of these microbiome-based interventions largely depends on the accurate identification of disease-associated microbial taxa [21]. Traditional abundance-based statistical analyses are limited to capturing complex interactions within the microbiome and potentially overlook critical disease-associated signals. To overcome this limitation, machine learning and deep learning-based models are increasingly being applied to learn complex patterns from high-dimensional microbiome data and uncover novel biomarkers relevant to disease prediction and treatment [22].

For example, Pasolli et al. [23] proposed MetAML, which applies random forests (RFs) and support vector machines (SVMs) to predict diseases using 2,424 publicly available microbiome samples. However, it only uses the abundance of the microbiome and does not incorporate microbial genomic information, such as DNA sequences, limiting its biological interpretability. Oh et al. [24] developed DeepMicro, an autoencoder-based deep learning framework that transforms microbial abundance data into low-dimensional representations, followed by classification using RF, SVM, and multilayer perceptron models for disease prediction. While effective in generating low-dimensional representations, this approach may result in information loss, and lacks end-to-end architecture. Liao et al. [25] introduced GDmicro, which constructs graphs in which each node represents an individual sample and latent features are derived for each node through domain adaptation (DA). These graphs were then used to predict diseases using graph convolutional networks. Although the DA enhances generalization across different cohorts, applying the model to new data requires graph reconstruction and retraining. Despite their methodological differences, existing models share common limitations: they often fail to incorporate microbial relationships and lack genomic information, which hinders both prediction performance and biological interpretability. Overcoming these limitations is complicated by the scarcity of publicly available gut microbiome data on AD. According to a recent study, only 220,017 human gut microbiome datasets are publicly available from the National Center for Biotechnology Information (NCBI) [26], and this number is likely to be substantially smaller when limited to AD.

To address these limitations, we propose ATOMIC, a graph attention neural network for ATOpic dermatitis prediction on human gut MICrobiome. ATOMIC is an interpretable deep learning model that incorporates microbial co-expression networks and microbial genomic information to predict AD. Unlike previous models that rely solely on abundance data, ATOMIC integrates the relationships among microbes and their genomic information into a graph-based architecture. By leveraging graph attention networks (GATs), the model learns representations that capture microbial relationships and identifies the key microbial contributors to AD prediction. Furthermore, it supports interpretability by highlighting microbe-level importance through attention scores, enabling the discovery of candidate biomarkers relevant to disease prediction and treatment. For development and evaluation, we applied ATOMIC to a new gut microbiome dataset collected from a cohort of adult patients with AD and released this dataset to the public.

## 2. Materials and methods

### 2.1 Overview of ATOMIC

An overview of ATOMIC is illustrated in Fig 1. We constructed a microbial co-expression network, where each node represented a microbe and each edge represented microbial co-expression. Microbes with zero-count abundance values were omitted from each sample to create sample-specific graphs, which led to variations in graph sizes between samples. The Graph Attention Network v2 (GATv2) [27] layers update the node representations by incorporating the relationships between the microbes and their neighboring nodes. Finally, the graph embedding is obtained through a self-attention readout, and AD prediction is performed using fully connected layers.

**Fig 1.**
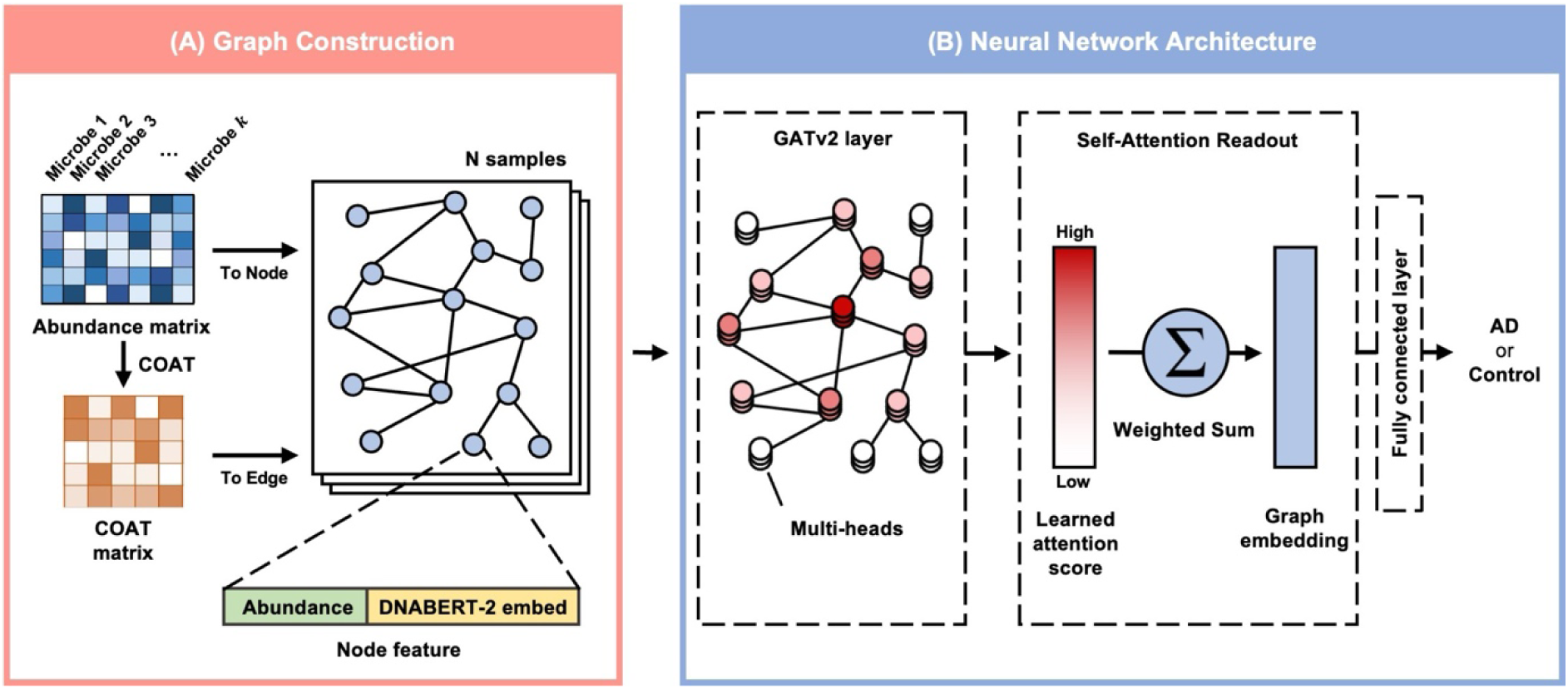
Overview of ATOMIC. (A) The microbial co-expression network construction process of ATOMIC. Each node represents a microbe with abundance and genomic vectors as features, while each edge represents a microbial co-expression. The microbial co-expression network is fed into GATv2 layers. (B) The neural network architecture of ATOMIC. Three stacked GATv2 layers with multi-head attention are followed by a self-attention readout to obtain the graph embedding, and fully connected layers use the graph embedding to predict AD. COAT, composition-adjusted thresholding; AD, atopic dermatitis.

### 2.2 Atopic dermatitis sample collection and data preprocessing

We collected gut microbiome data from 99 adult participants recruited through the Department of Dermatology at Kangwon National University Hospital (KNUH) in Chuncheon, Korea. The cohort included 70 patients diagnosed with AD, aged 18–69 years based on the Hanifin and Rajka criteria, and 29 healthy controls without chronic inflammatory or autoimmune diseases. Participants were excluded if they had used systemic immunosuppressants or steroids, had a history of inflammatory or autoimmune diseases, or were unable to visit within ±1 week of the scheduled date. The study protocol was approved by the Institutional Review Board of KNUH (No. KNUH-2023-08-011-002) and written informed consent was obtained from all participants. Stool samples were collected in sterile containers and stored at –80 °C within 4 hours of collection. All collected samples were stored in ultra-low temperature freezers (–80 °C or below) or liquid nitrogen tanks (–130 °C to –196 °C) in the human biobank. After completion of the analyses, the remaining samples were disposed according to the disposal protocols of the analysis institution.

To augment the sample size and improve the generalizability of the microbial co-expression network construction, we additionally collected 1,392 samples from patients with AD and healthy controls available from the NCBI database. These publicly available datasets served as external resources for training more robust models by enhancing microbial association inferences. These additional samples were derived from five independent studies conducted in Korea [28], Hong Kong [29], China [30, 31], and Japan [32]. An overview of all the collected samples is summarized in Table 1.

**Table 1.**
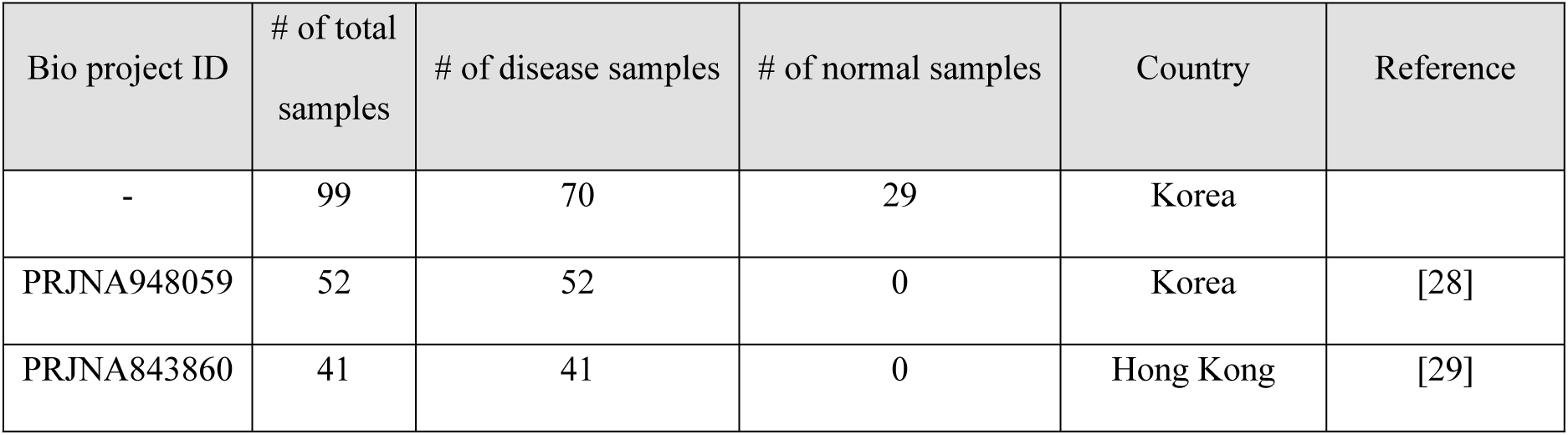
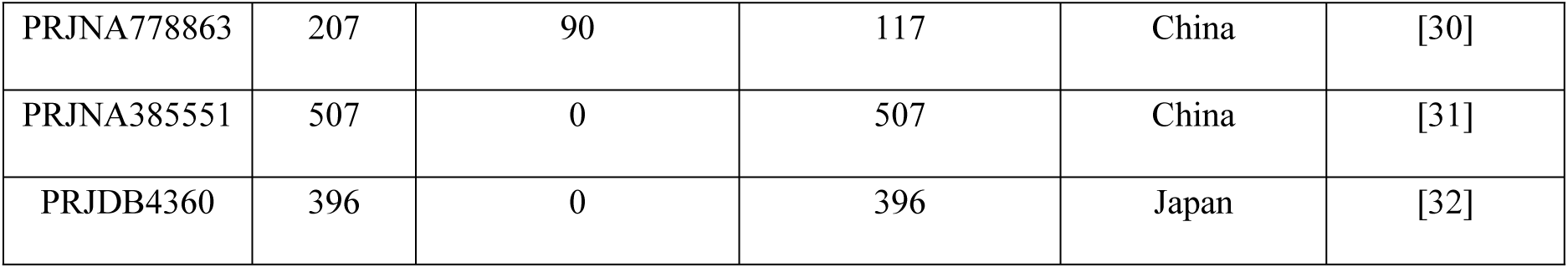
Detailed information about the collected data used in this study.

All sequencing data, including those from KNUH and public repositories, were sequenced using Illumina paired-end sequencing platform targeting 16S rRNA amplicon sequencing and they were processed using a unified bioinformatics pipeline. We performed microbial taxonomic classification and abundance estimation based on the SILVA 138.1 reference database [33] after preprocessing all sequences using QIIME2. However, owing to the low proportion of sequences matching at the species level, we aggregated the abundance of species to obtain genus-level microbial abundance data. Genus-level abundances were then normalized within each sample to a sum of 100.

### 2.3 Microbial co-expression network construction

We constructed a global microbial co-expression network as an undirected graph 𝐺 = (𝑉,𝐸), where the nodes represent microbes and the edges indicate statistically significant co-abundance relationships. To define the edges, we computed correlation coefficients between microbial taxa using the composition-adjusted thresholding (COAT) method [34], which infers correlations based on the log ratios between pairs of compositional variables. To construct a robust network, we randomly sampled 80% of the dataset five times and computed COAT correlations in each iteration. Only microbial correlations consistently observed across all five iterations were retained as edges in the final network. Note that the microbial co-expression network was constructed using both the KNUH and public datasets, while model training and evaluation were performed exclusively on the KNUH cohort.

We represent the set of 𝑛 microbial nodes as 𝑉 = {(𝑣_𝑖_, 𝑠_𝑖_)}^𝑛^ where each node 𝑖 is characterized by a 64- dimensional abundance vector 𝑣_𝑖_ ∈ ℝ^64^ and a 768-dimensional genomic vector 𝑠_𝑖_ ∈ ℝ^768^. The abundance vector 𝑣_𝑖_ for microbe 𝑖 was obtained by multiplying the scalar abundance of microbe 𝑖 with a 64-dimensional learnable vector that was randomly initialized from a uniform distribution. The genomic vector 𝑠_𝑖_ was derived from DNABERT-2 [35], a genome foundation model. Specifically, for each genus-level taxon, we encoded the DNA sequences of all available constituent species and then computed the average of these vectors. Thus, each microbial node 𝑖 has an initial representation ℎ^(0)^ ∈ 𝑅^832^, obtained by concatenating its abundance vector 𝑣_𝑖_ and genomic vector 𝑠_𝑖_. The connections between these nodes are defined by the edge set 𝐸 = {𝑒_𝑖𝑗_||𝑐_𝑖𝑗_| ≥ 0.1, 𝑖,𝑗 = 1,2,…,𝑛, 𝑖 ≠ 𝑗}, where 𝑒_𝑖𝑗_ represents a co-expression edge between microbial nodes (𝑣_𝑖_, 𝑠_𝑖_) and (𝑣_𝑗_, 𝑠_𝑗_), and 𝑐_𝑖𝑗_ is their corresponding COAT correlation coefficient.

Although the topology of the co-expression graph (i.e., edge set 𝐸) was fixed across all samples, the set of active nodes varied depending on the sample-specific microbial composition. For each sample, we constructed a subgraph by removing nodes corresponding to microbes with zero abundance. As a result, each sample had a distinct graph size, reflecting its individual microbial profile. This strategy enables sample-specific graph representation while preserving a consistent co-expression backbone, facilitating efficient learning of microbial interactions in a biologically meaningful and scalable manner.

### 2.4 Graph neural network for learning microbial co-expression relationships

After constructing the microbial co-expression networks, we applied GATv2 to learn relationships among microbes and predict AD. GATv2 is an improved architecture over the original GAT [36] that addresses the limitation of static attention, where certain key nodes consistently receive high attention weights regardless of the query node. This static ranking reduced the capacity of the model to capture context-dependent interactions. By contrast, GATv2 introduces dynamic attention, allowing attention weights to vary depending on the query node. This enables a more expressive and flexible modeling of complex relational structures in the graph, which is particularly important for capturing sample-specific microbial interactions.

In each GATv2 layer 𝑙, the importance of a neighboring node 𝑗 to a target node 𝑖 is computed using the attention coefficients. GATv2 adopts a multi-head attention mechanism in which each attention head *k* independently learns distinct attention patterns. These attention coefficients are normalized across all neighbors 𝑗 ∈ 𝑁_𝑖_ using softmax function [37], formally defined as:

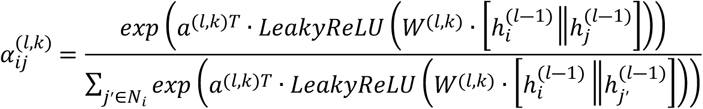

where 𝑎^(𝑙,𝑘)^ denotes the attention vector, 𝑊^(𝑙,𝑘)^ denotes the linear transformation matrix at layer 𝑙 for the 𝑘-th attention head, and || denotes the vector concatenation.

The updated representation of node 𝑖 in layer 𝑙 is computed by aggregating its neighbor’s features weighted by the attention coefficients across all 𝐾 heads:

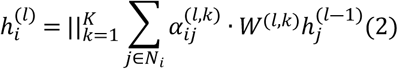

where ℎ^(𝑙)^ denotes the updated representation of node 𝑖 in layer 𝑙, 𝐾 is the number of attention heads; and 𝑊^(𝑙,𝑘)^ is the linear transformation matrix corresponding 𝑘-th 𝑙attention head in layer 𝑙.

In the final GATv2 layer 𝐿, the node embeddings from all the attention heads are averaged:

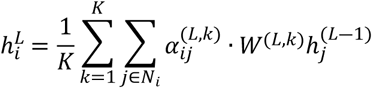

The resulting node embeddings ℎ^𝐿^ = [ℎ^𝐿^_1_,ℎ^𝐿^_2_,…,ℎ^𝐿^_𝑛_], ℎ^𝐿^_𝑖_ ∈ ℝ^𝐹^, where 𝐹 denotes the dimensionality of the updated node embeddings, are used to compute the graph embedding through an attention-based readout. Each node embedding ℎ^𝐿^_𝑖_ is projected to a scalar importance score, ℎ_𝑖_, using a learnable linear transformation matrix

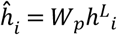

To incorporate the attention-based node importance into the graph embedding for the prediction of AD, we calculated the attention score 𝐴_𝑖_ for each node 𝑖 by applying the softmax function as:

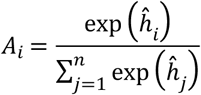

The final graph embedding 𝑃 is then computed as the weighted sum of the node embeddings using the attention score 𝐴_𝑖_. Finally, the graph embedding 𝑃 is fed into a fully connected network with two hidden layers to predict the AD. In our implementation, we used three stacked GATv2 layers (L = 3), each configured with eight attention heads (k = 8). The output dimensions of each node representation were set to 32 (F = 32). To improve generalization, we applied edge dropout (p = 0.3), Mish activation [38], node dropout (p = 0.3) [39].

### 2.5 Model Implementation

Our dataset of 99 samples was split into a training set of 59 samples, a validation set of 20 samples, and a test set of 20 samples, corresponding to an approximate 60:20:20 ratio. We implemented ATOMIC neural network architecture using PyTorch [40] and optimized the model using the AdamW optimizer [41]. The initial learning rate was set as 0.0001, with a batch size of 16, and the learning rate was scheduled to decay by 1% every 10 epochs. The hyperparameters, including the number of layers, number of heads, dropout rates, and learning rate, were optimized based on the grid search using a 4-fold cross-validation strategy to minimize validation loss. We trained our model on a computing machine equipped with an Intel Xeon Gold 6230 CPU, 512 GB Memory, and an NVIDIA A100 GPU.

## 3. Results

### 3.1 Performance on the KNUH dataset

We evaluated the performance of ATOMIC on the KNUH dataset by comparing it with several baseline models, including deep learning-based methods such as GDmicro, DeepMicro, and MetAML, as well as traditional machine learning models such as RFs and SVMs (Table 2). GDmicro was originally designed with DA to enhance cross-cohort generalization. However, because our model focused on optimizing performance within a single AD cohort, we disabled the DA component of GDmicro to ensure a fair and direct comparison within the same cohort setting.

**Table 2.** Performance comparison of baseline models on the KNUH dataset in terms of AUROC, AUPRC, and F1 score.

| Model | AUROC | AUPRC | F1 |
| --- | --- | --- | --- |
| RFs | 0.595 (0.060) | 0.782 (0.048) | 0.622 (0.224) |
| SVMs | 0.536 (0.000) | 0.785 (0.000) | 0.444 (0.000) |
| MetAML | 0.548 (0.033) | 0.706 (0.024) | 0.769 (0.024) |
| DeepMicro | 0.579 (0.032) | 0.768 (0.017) | <u>0.835 (0.034)</u> |
| GDmicro (w/o DA) | 0.805 (0.029) | 0.664 (0.118) | 0.742 (0.052) |
| GDmicro (w/o DA, ensemble) | **0.893** | 0.788 | 0.750 |
| ATOMIC | 0.752 (0.069) | <u>0.894 (0.027)</u> | 0.784 (0.108) |
| ATOMIC (ensemble) | <u>0.810</u> | **0.927** | **0.846** |

As a result, ATOMIC outperformed most baseline models, achieving an area under the receiver operating characteristic curve (AUROC) of 0.752 ± 0.069, area under the precision-recall curve (AUPRC) of 0.894 ± 0.027 and an F1 score of 0.784 ± 0.108 over five independent runs with different random seeds. Although some performance variability was observed, likely owing to the limited training sample size, the model demonstrated a strong overall predictive capability. Incorporating ensemble learning further improved both stability and performance. Specifically, the ensemble-based ATOMIC boosted the AUROC by 7.71%, AUPRC by 3.69%, and F1 score by 7.91% compared with the non-ensemble version. Compared with GDmicro without DA, our ensemble-based ATOMIC achieved competitive AUROC performance and showed substantial gains in AUPRC (17.6%) and F1 score (12.8%), indicating greater robustness and discriminative power. Notably, GDmicro without DA exhibits a higher AUPRC variance, whereas our model maintains a more stable performance across runs. Statistical significance was assessed using the Mann–Whitney U test. ATOMIC with the ensemble achieved significantly higher AUROC scores than the RFs (p = 0.004), SVMs (p = 0.002), MetAML (p = 0.004), and DeepMicro (p = 0.004), confirming the robustness of our approach for identifying true positives. Some baseline models showed relatively lower AUROC performance, possibly because of their limited capacity to capture nonlinear inter-microbial dependencies and a lack of microbial genomic feature integration. In contrast, ATOMIC benefits from leveraging microbial co-expression patterns, enabling more effective modeling of complex microbial interactions, and ultimately leading to more accurate and stable predictions of AD.

Boldface indicates the best performance, underlining denotes the second-best, and all values are reported as mean (standard deviation) across five independent runs. Abbreviations: KNUH, Kangwon National University Hospital; AUROC, area under the receiver operating characteristic curve; AUPRC, area under the precision-recall curve; RFs, random forests; SVMs, support vector machines; DA, domain adaptation.

We visualized sample representations using both graph embeddings derived from the self-attention readout and raw microbial abundance data to examine the clustering patterns between the AD and healthy control samples. Fig 2 shows the t-SNE plots on the KNUH dataset, where panel A corresponds to graph embeddings and panel B corresponds to microbial abundance profiles. The graph-based representations revealed more distinct cluster boundaries between the AD and control groups compared to the abundance-based features. To quantitatively assess group separability, we calculated silhouette scores for both representations. The graph embeddings yielded a significantly higher silhouette score (0.566, p = 0.001) than the microbial abundance data (0.055, p = 0.119), suggesting superior class separation. These results demonstrate that the graph-based attention mechanism of ATOMIC captures latent structural patterns in the microbiome that are not apparent from raw abundance alone, thereby contributing to its enhanced classification performance.

**Fig 2.**
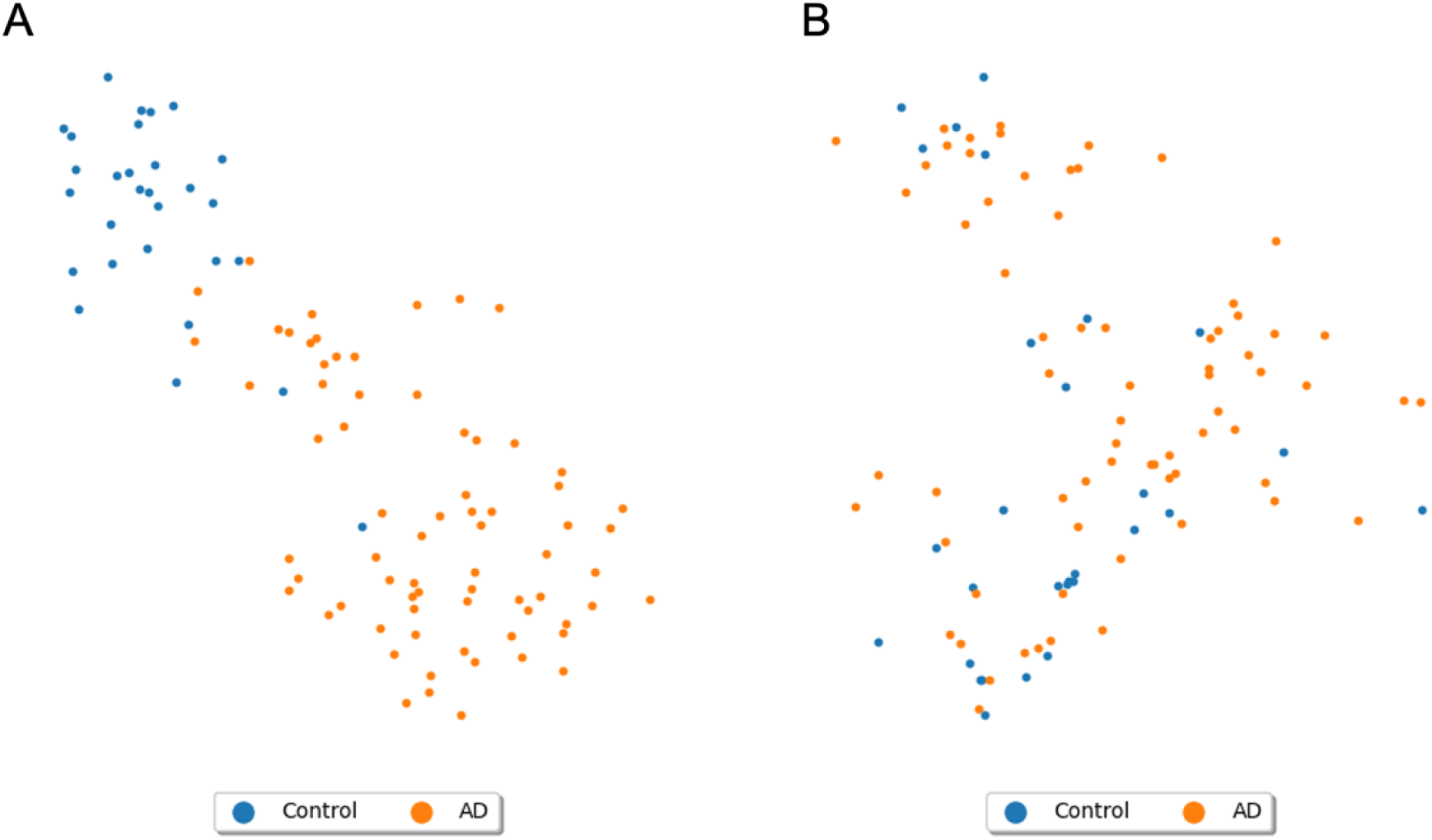
t-SNE visualization of the KNUH dataset (orange: AD; blue: healthy control). (A) Graph embeddings derived from ATOMIC. (B) Genus-level microbial abundance data. KNUH, Kangwon National University Hospital; AD, atopic dermatitis.

### 3.2 Ablation study

To assess the contribution of individual components within ATOMIC, we conducted ablation experiments by removing the microbial genomic information and altering the structure of the co-expression network. As shown in Table 3, excluding genomic features from node representations led to a significant decline in model performance (p = 0.045, Mann–Whitney U test), highlighting the importance of integrating genomic context alongside abundance data for microbial feature characterization. Furthermore, when the genomic information was removed and the edges of the co-expression graph were randomly shuffled for each sample, the performance declined even further (p = 0.028). This indicates that the structured microbial relationships encoded in the co-expression network play a key role in enhancing the predictive accuracy. Collectively, these results confirm that both microbial genomic features and biologically meaningful co-expression structures are essential for ATOMIC’s robust prediction of AD.

**Table 3.** Ablation study evaluating the impact of microbial genomic features and co-expression graph structure on ATOMIC performance.

| Model | AUROC | AUPRC | F1 |
| --- | --- | --- | --- |
| ATOMIC | 0.752 (0.069) | 0.894 (0.027) | 0.784 (0.108) |
| w/o genomic information | 0.674 (0.065) | 0.851 (0.028) | 0.680 (0.144) |
| w/o genomic information +<br>edge shuffle | 0.657 (0.046) | 0.839 (0.035) | 0.495 (0.292) |

All values are reported as mean (standard deviation) across five independent runs. Abbreviations: AUROC, area under the receiver operating characteristic curve; AUPRC, area under the precision-recall curve; w/o, without.

### 3.3 Model interpretation with attention scores

A key advantage of ATOMIC is its interpretability, which is enabled by the attention scores derived from the self-attention readout layer. In the graph representation, each node corresponds to a microbial genus, and the attention score 𝐴_𝑖_ for node 𝑖 quantifies its relative importance in the model’s prediction. Since the attention scores across all nodes in a sample sum to 1, they allow for within-sample comparisons of microbial relevance. Note that these scores are non-directional; a high score indicates predictive importance but does not imply whether the microbe is enriched or depleted in that sample.

To identify microbes most associated with AD predictions, we aggregated attention scores across samples correctly classified as AD in the test set. Since each sample contained a different subset of microbes, a direct comparison of raw attention scores across samples was inappropriate. To address this issue, we applied a centered log-ratio (CLR) transformation, which is a common approach for compositional data normalization. CLR transforms each attention score by dividing it by the geometric mean of all scores in the same sample, enabling inter-sample comparisons. After normalization, we calculated the average CLR-transformed attention scores for each microbe and ranked them. Table 4 lists the top 10 genera with the highest CLR-transformed attention scores, along with the supporting literature, all of which have been previously implicated in AD. Because high attention scores reflect predictive contributions, both protective and risk-associated taxa can be prioritized if they provide informative patterns for AD classification.

**Table 4.** Literature-supported summary of the top 10 microbial genera ranked by CLR-transformed attention scores from the trained ATOMIC model.

| Rank | Genus | CLR-normed Attention Score | Literature |
| --- | --- | --- | --- |
| 1 | *Ruminococcus_gnavus_group* | 4.877 | [42] |
| 2 | *Dorea* | 4.827 | [50] |
| 3 | *Butyricoccus* | 4.824 | [45] |
| 4 | *Flavonifractor*<br>(family *Oscillospiraceae*) | 4.800 | [50] |
| 5 | *Blautia* | 4.772 | [52] |
| 6 | *Streptococcus* | 4.767 | [53] |
| 7 | *Coprococcus* | 4.703 | [46] |
| 8 | *Lachnoclostridium* | 4.515 | [47] |
| 9 | *Rothia* (family *Micrococcaceae*) | 4.091 | [50] |
| 10 | *Eubacterium_hallii_group* | 4.045 | [48] |

Abbreviations: CLR, centered log-ratio.

The highest-ranked genus was *Ruminococcus_gnavus_group* (CLR score = 4.877), a well-known producer of SCFAs [42]. SCFAs promote the differentiation of regulatory T cells (Tregs) [43], support intestinal and epidermal barrier integrity, and exert broad anti-inflammatory effects [44]. Other highly ranked genera, including *Butyricicoccus*, *Coprococcus*, *Lachnoclostridium*, and *Eubacterium_hallii_group*—are also recognized producers of SCFAs, especially butyrate [45–48]. Notably, reduced SCFA levels have been consistently observed in children with AD [49], further supporting their protective role. Additionally, *Dorea* (family *Lachnospiraceae*), *Flavonifractor* (family *Oscillospiraceae*), and *Rothia* (family *Micrococcaceae*) have been reported to decrease or negatively correlate with AD severity [50, 51].

In contrast, ATOMIC also identified genera associated with increased risk of AD. *Blautia* and *Streptococcus* both received high attention scores and have been shown to be more abundant in individuals with AD compared to healthy controls [52, 53]. Notably, *Streptococcus* has been linked to persistent forms of AD rather than transient presentations [53], suggesting a role in chronic inflammation. These findings demonstrate that ATOMIC is not only effective in predicting AD, but also in suggesting biologically relevant microbial patterns. Its attention-based mechanism provides interpretable outputs that align with known disease mechanisms, offering valuable insights into microbial contributions to AD pathogenesis.

### 3.4 Biomarker discovery from microbe-mediated genes

We further analyzed the potential functions of microbes that with high attention scores captured by ATOMIC. To accomplish this, we conducted microbe function analysis using Microbe-Set Enrichment Analysis (MSEA) [54]. We utilized a publicly available microbe-set library linking human genes to genus-level microbes and applied a custom background consisting of microbial genera detected in the KNUH dataset. To analyze only the microbe sets that primarily contributed to AD prediction, we selected microbes with average CLR-normalized attention scores greater than zero, calculated from AD test samples, as input for MSEA. The resulting enriched genes with an adjusted p-value below 0.05 are shown in Fig 3.

**Fig 3.**
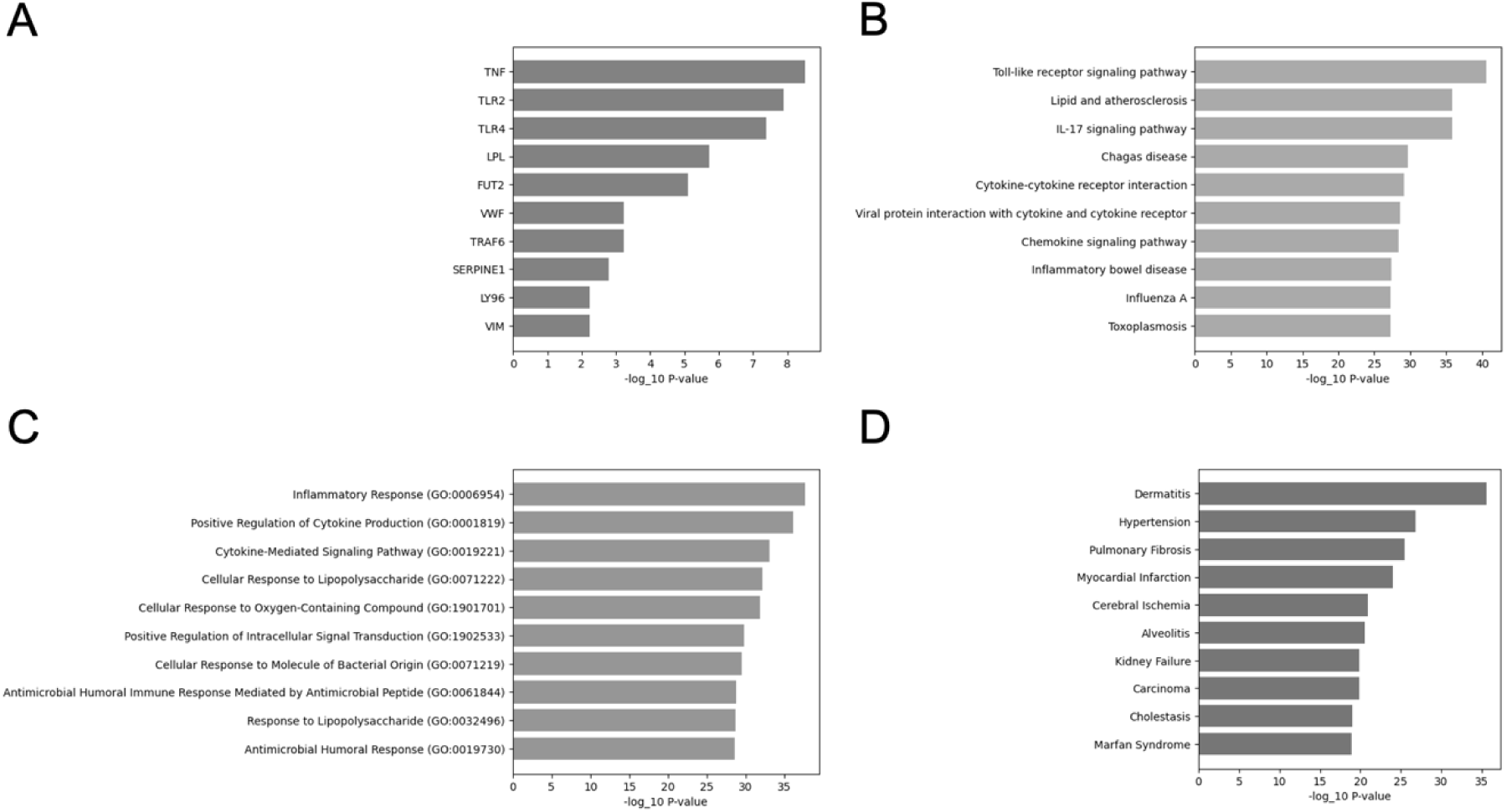
Bar plots illustrating enrichment analysis results. (A) Top 10 enriched microbe-mediated genes identified by Microbe-Set Enrichment Analysis (MSEA). (B) Top 10 enriched pathway terms from the KEGG 2021 Human database. (C) Top 10 enriched Gene Ontology terms from the GO Biological Process 2025. (D) Top 10 enriched disease ontology terms from GeDiPNet 2023. KEGG, Kyoto Encyclopedia of Genes and Genomes; GO, Gene Ontology.

Fig 3A shows the top 10 enriched human genes functionally linked to microbes predictive of AD. Many of these genes are key regulators of Th2-mediated inflammation, which is a hallmark of AD pathogenesis. For example, TNF encodes TNF-α, a key cytokine elevated in the skin of AD patients [55]. TNF-α promotes the expression of Th2 cytokines and TSLP [56], both of which aggravate Th2-mediated immune responses by activating dendritic and mast cells [57].

We also identified innate immune receptors such as Toll-like receptor (TLR)-2 and TLR4 among the top-ranked genes. TLR2 promotes Th2 polarization through IL-5 and FcεRI upregulation, while TLR4 broadly modulates inflammatory signaling [58, 59]. Polymorphisms in both TLR2 and TLR4 have been associated with increased susceptibility to AD [60]. These receptors are also functionally linked to the microbial genera identified by ATOMIC, such as *Butyricicoccus*, *Coprococcus,* and *Ruminococcus_gnavus*, which produce SCFAs and modulate host immunity through TLR4 activation. SCFAs are known to suppress inflammatory cytokines including TNF-α, IL-6, and IL-8, yet SCFA level are reduced in patients with AD [61]. Moreover, *Ruminococcus_gnavus* secretes glucorhamnan, which stimulates dendritic cells to produce TNF-α through the TLR4 signaling pathway [62], directly linking this microbe to inflammation-relevant host responses.

Several other inflammation-related genes have been identified in addition to canonical immune mediators. LY96, which encodes the MD-2 protein, functions as an essential coreceptor in the TLR4 signaling pathway and has been proposed as a target for genetic modulation in AD [63]. TRAF6 is another key immune regulator previously implicated in IL-23/IL-17-mediated responses in psoriasis-like dermatitis [64], and may play a parallel role in AD.

Genes associated with metabolic dysfunction in AD were also identified in AD. LPL is particularly notable, given the alterations in lipid metabolism observed in patients with AD; LPL agonists are currently under investigation as potential therapeutic agents [65–67]. Similarly, SERPINE1 has been reported to be significantly upregulated in AD lesions and may contribute to dermatitis-associated pruritus [68].

Importantly, our analysis identified several genes that have not been previously associated with AD, highlighting potential novel biomarker candidates. For example, VWF, although not directly linked to AD, is known to activate macrophages and stimulate the production of pro-inflammatory mediators such as TNF, IL-6, CCL2, and CCL3 [69]. Similarly, FUT2 and VIM have been implicated in other chronic inflammatory conditions, such as inflammatory bowel disease, but their roles in AD remain uncharacterized [70, 71]. Nevertheless, given the critical role of inflammatory responses in AD pathogenesis [72], these genes warrant further investigation as potential biomarkers or therapeutic targets for AD.

### 3.5 Function and disease annotation of microbe-mediated genes

To investigate the biological functions of microbe-mediated genes identified by ATOMIC, we conducted a secondary enrichment analysis using Enrichr [73]. Specifically, we selected significantly enriched gene sets from the prior MSEA results (adjusted p < 0.05) and analyzed them across pathways, gene ontology, and disease annotation databases. Pathway enrichment analysis using the KEGG 2021 human database revealed several immune-related pathways that are implicated in AD pathogenesis (Fig 3B). Notably, the toll-like receptor (TLR) signaling pathway (rank 1) was highly enriched. TLRs are pattern recognition receptors essential for microbial sensing and innate immune activation [74]. TLR agonists have been shown to suppress Th2-mediated responses in CD4+ T cells, offering potential therapeutic benefits in AD [75]. The IL-17 signaling pathway (rank 3) also appeared, consistent with its known role in exacerbating AD through the suppression of FLG expression, which is vital for epidermal barrier integrity [76]. Cytokine–cytokine receptor interactions (rank 5) and chemokine signaling pathways (rank 7) were similarly enriched. These pathways include cytokines, such as IL-4, IL-13, IL- 31, and TSLP, and chemokines, such as CCL1, CCL13, and CCL17, which have been implicated in barrier disruption, pruritus, and immune cell recruitment in AD [77, 78].

Gene Ontology analysis (GO Biological Process 2025; Fig 3C) further confirmed the enrichment of the immune and inflammatory processes. Terms such as “inflammatory response” (GO:0006954), “positive regulation of cytokine production” (GO:0001819), and “cytokine-mediated signaling pathway” (GO:0019221) were highly ranked, aligning with both literature-based expectations and KEGG pathway results. Disease ontology enrichment using GeDiPNet 2023 (Fig 3D) also highlighted “dermatitis” as the most significantly associated disease term, reinforcing the AD relevance of the gene sets prioritized by ATOMIC. Collectively, these results demonstrate that ATOMIC not only prioritizes predictive microbial genera but also recovers host gene signatures and immune pathways highly relevant to AD pathophysiology, supporting its potential utility for both diagnostic and mechanistic insights.

### 3.6 Microbial composition analysis

To explore the therapeutic implications of ATOMIC’s microbe-level attention scores, we designed an *in silico* intervention experiment to simulate the effect of removing specific microbial taxa. This analysis aimed to evaluate whether the microbes identified as influential by ATOMIC contributed meaningfully to AD predictions and could serve as potential targets for microbiome-based treatment strategies in AD.

We stratified microbial genera into two groups based on their average CLR-normalized attention scores in the AD-classified test samples: Group A included taxa with positive attention scores, representing microbes that positively contributed to AD predictions, whereas Group B included taxa with negative attention scores, presumed to have minimal influence. To simulate therapeutic removal, we randomly eliminated 30%, 50%, and 70% of the microbes from each group in the AD test samples and measured the resulting changes in the predicted AD probabilities. If Group A microbes are truly predictive and causally linked to disease severity, their removal should reduce the model’s confidence in AD classification. Conversely, the removal of microbes from Group B was expected to have little to no effect. Each removal scenario is repeated 100 times to ensure statistical robustness. The results are summarized in Table 5.

**Table 5.**
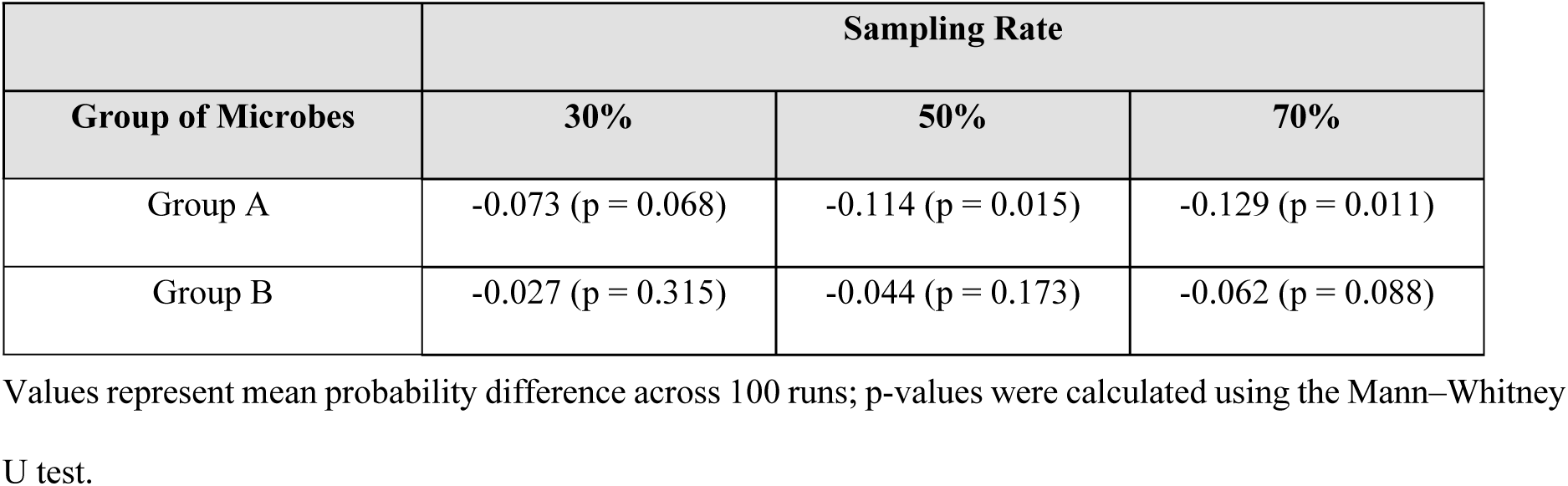
Effects of alterations in microbial composition on the prediction of AD probability in ATOMIC.

|  | Sampling Rate |  |  |
| --- | --- | --- | --- |
| Group of Microbes | 30% | 50% | 70% |
| Group A | -0.073 (p = 0.068) | -0.114 (p = 0.015) | -0.129 (p = 0.011) |
| Group B | -0.027 (p = 0.315) | -0.044 (p = 0.173) | -0.062 (p = 0.088) |

Values represent mean probability difference across 100 runs; p-values were calculated using the Mann–Whitney U test.

Removal of Group A taxa led to a progressive decrease in the predicted AD probabilities, with reductions ranging from 0.075 to 0.125, depending on the removal fraction. These decreases were statistically significant at 50% (p = 0.015) and 70% (p = 0.011) removal levels, as determined by the Mann–Whitney U test. In contrast, the removal of Group B taxa had minimal impact across all sampling rates.

These findings support the hypothesis that ATOMIC effectively identifies microbial taxa that substantially influence AD prediction. This *in silico* intervention framework provides a useful approach for prioritizing microbes for further investigation of their functional roles in AD. While this does not imply that removing these microbes would directly confer therapeutic benefits, it suggests that high-attention taxa may represent ecologically or immunologically important members of the AD-associated microbiota.

## 4. Discussion

This study demonstrated that ATOMIC outperforms baseline models for prediction of AD and provides interpretability. Its superior performance in classifying AD results from the incorporation of microbial co-expression networks and microbial genomic information, enabling ATOMIC to capture both structural and functional relationships within the microbiome. ATOMIC assigns microbe-specific attention scores, thereby capturing the relative importance of each microbe. Additionally, to promote broader research on the relationship between atopic dermatitis and the gut microbiome, we made the processed abundance data from this study publicly available to the research community.

Through a literature-based analysis of high-attention microbes, we identified associations between known AD-related mechanisms, including SCFA production and host immune modulation. Furthermore, enrichment analyses of microbe-mediated genes revealed significant involvement in inflammation-related pathways, such as TLR signaling, IL-17 signaling, and cytokine–cytokine receptor interactions. Interestingly, genes, such as VWF, FUT2, and VIM, which are not directly linked to AD, have been implicated in broader inflammatory conditions, suggesting novel avenues for future research.

Beyond prediction, ATOMIC provides a framework for individualized microbiome interpretation. Given the known inter-individual variability in the gut microbial composition, patients with similar clinical presentations may respond differently to treatment. ATOMIC addresses this challenge by identifying the most influential microbes in each individual, thereby supporting personalized diagnostic and therapeutic strategies. Its attention mechanism highlights taxa most predictive for each individual, offering interpretable insights that can inform clinical decision-making. While conventional probiotics and FMT aim to restore the overall microbial balance, their broad-spectrum and non-specific nature limits their utility for disease-specific interventions [79]. These findings the need for next-generation probiotics (NGPs), which involve targeted modulation of disease-associated taxa [79, 80]. In this context, ATOMIC’s attention scores offer a data-driven approach to prioritize candidate NGP strains for AD, enabling precision targeting on a per-patient basis, bridging the gap between microbiome analysis and personalized medicine in AD.

Despite these contributions, several limitations should be acknowledged. First, we used genus-level rather than species-level data, potentially obscuring species-specific effects. Second, while our training data consisted solely of adult samples, some supporting literature referenced pediatric studies, introducing possible age-related confounders [81]. Nevertheless, specific treatment guidelines for adult-onset AD have not yet been established; consequently, therapeutic approaches developed for pediatric AD are commonly applied in adults [82]. The findings of this study may serve as a valuable reference for future studies aimed at developing targeted therapeutic agents for adult-onset AD. Third, the KNUH dataset is relatively small (n = 99), which may have limited the generalizability of our findings. Although cross-validation was employed to reduce overfitting, external validation was essential.

To address this, we applied ATOMIC to an external pediatric AD dataset from Korea [83]. This dataset was obtained from the European Nucleotide Archive (accession No. PRJEB41351) comprised 346 samples, including 234 patients with AD and 112 healthy controls. On this dataset, the ATOMIC ensemble achieved an AUPRC of 0.886 and an F1-score of 0.762, surpassing GDmicro-AUPRC (0.773) and F1 (0.733), although GDmicro yielded a higher AUROC of 0.867 compared to 0.735 for ATOMIC. This result was consistent with the pattern observed in the KNUH dataset. Although the AUROC was relatively low, ATOMIC maintained a higher AUPRC and F1 score, suggesting better precision in identifying true AD cases, which may be advantageous for clinical applications that prioritize specificity. Interestingly, the performance of ATOMIC decreased on the pediatric dataset, despite the larger sample size. This performance gap is likely due to age-dependent differences in the gut microbiome composition [84], which in turn limits the applicability of adult-based microbial co-expression networks to pediatric data.

Future studies will focus on improving the generalizability of the model by incorporating diverse microbiome datasets across age groups, geographic regions, and disease subtypes. In addition, we plan to enhance the scalability of ATOMIC by integrating techniques such as deep domain adaptation [85] and knowledge distillation [86], enabling effective transfer to large-scale and heterogeneous datasets.

## 5. Conclusion

In this study, we presented ATOMIC, an interpretable graph attention network-based model that integrates microbial co-expression networks with genomic information to predict AD using gut microbiome data. By combining genus-level abundance profiles and DNABERT-derived genomic features as graph node attributes, ATOMIC effectively captures both structural and functional relationships among microbes. Trained on microbiome data from adult patients with AD and healthy controls, ATOMIC outperformed the existing baseline models in predictive performance. Importantly, the attention scores derived from the model enabled the identification of key microbial taxa that contributed to the AD classification. Functional enrichment analysis of microbe-mediated genes revealed host immune pathways and inflammation-related mechanisms associated with these microbes. These findings highlight ATOMIC’s potential as a predictive model and tool for discovering candidate microbial biomarkers and therapeutic targets for AD. To support continued research, we publicly released the processed gut microbiome data. ATOMIC offers a foundation for future efforts toward personalized microbiome-based interventions and biomarker discovery for AD.

## Supporting information

Not applicable.

## Acknowledgments

This study was supported by the Ministry of Science and ICT (MSIT), Korea, under the Technology Development Program (S3364091) of the MSS, the ICT Challenge and Advanced Network of HRD (ICAN) program (IITP- 2025-RS2022-00156439) supervised by the Institute of Information & Communications Technology Planning & Evaluation (IITP), and the Bio&Medical Technology Development Program of the National Research Foundation (NRF) funded by the Korean government (MSIT) (No. RS-2024-00441029).

## Data availability statement

The source code and processed abundance data are available from ATOMIC GitHub Repository at https://www.github.com/KU-MedAI/ATOMIC. The raw sequence data are available upon request.

